# Modeling the effects of genetic and diet induced obesity on melanoma progression in zebrafish

**DOI:** 10.1101/2022.05.27.493792

**Authors:** Emily Montal, Dianne Lumaquin, Yilun Ma, Shruthy Suresh, Richard M. White

## Abstract

Obesity is a rising concern and associated with an increase in numerous cancers often in a sex-specific manner. Preclinical models are needed to deconvolute the intersection between obesity, sex, and cancer. We have generated a zebrafish system that can be used as a platform for studying these factors. We studied how germline overexpression of AgRP along with a high-fat diet (HFD) affects melanomas dependent on BRAF^V600E^. This revealed an increase in tumor incidence and area in male obese fish, but not females, consistent with the clinical literature. This is dependent on the somatic mutations, as male tumors generated with an RB1 mutation are sensitive to obesity, but this is not observed with PTEN. These data indicate that both germline and somatic mutations contribute to obesity related effects in melanoma. Given the rapid genetic tools available in the zebrafish, this provides a high-throughput system to dissect the interactions of genetics, diet, sex, and host factors in obesity-related cancers.

**Summary Statement:** Due to the rising incidence of obesity, there is a corresponding increased occurrence of obesity related cancers, which is often described to be dependent on sex. Here we developed a model to investigate the intersection between obesity, sex, and cancer.

## Introduction

There has been a sharp rise in the number of overweight and obese people in the US and worldwide. The Centers for Disease Control and Prevention reported in 2018 that 42% of Americans over the age of 20 are obese or severely obese, an increase from 30% in 2000 (Hales et al., 2020). As the incidence of obesity increases, so does the prevalence of diseases that are associated with obesity. This includes heart disease, type 2 diabetes, metabolic syndrome, and several cancers, including breast, colorectal, pancreatic cancer and melanoma (Renehan et al., 2008; Wolk et al., 2001).

Melanoma is the most lethal form of skin cancer. It is derived from melanocytes or their neural crest precursors in the skin. As melanoma invades deeper into the skin, the cells encounter several microenvironmental cell types including adipocytes. These adipocytes contribute to melanoma invasiveness by providing fatty acids as fuel for growth (Zhang et al., 2018). Obesity is well known to increase adipocyte numbers and sizes, and thus it is likely that the cross-talk between melanoma and adipocytes would be enhanced in obese individuals (Björntorp and Sjöström, 1971; Verboven et al., 2018). Clinically, the effect of obesity on melanoma is suggested but the data is not entirely clear. For example, clinical studies suggest that there is an increased hazard ratio for melanoma associated with obesity, but this is only in males (Karimi et al., 2016; Sergentanis et al., 2013). Further adding to this complexity, targeted therapy has been shown to be more effective in the obese setting only in males, suggesting a sex dependent effect of obesity on melanoma (McQuade et al., 2018). Preclinical animal studies using both genetic and diet models of obesity in mice demonstrate that there is an increase in xenograft tumor size in the obese setting (Brandon et al., 2009; Pandey et al., 2012; Ringel et al., 2020). These studies however do not address the effect of sex in these animals, nor do they test the effect of obesity on response to therapy. Furthermore, studies focusing on the systemic versus local effect of obesity on melanoma have been limited. Therefore, there is a critical need to develop preclinical models to further clarify the questions surrounding obesity, sex and melanoma.

Because the effects of obesity on cancer occur within the context of host physiology, it is important to study this in model organisms. The mouse is the most traditional model for this work, given its amenability to genetic manipulation, robust cancer models and ease of dietary interventions. However, there are limitations to mice in terms of the number of genetic manipulations that can be done, often requiring large numbers of crosses to gain germline alleles, and challenges in performing *in vivo* unbiased screens. In addition, detailed *in vivo* imaging remains difficult in the mouse outside of specialized equipment.

The zebrafish has emerged as an important model organism in cancer biology. The major advantage of the model is that it is highly amenable to large-scale and rapid genetic manipulation (using CRISPR or cDNA screens), allows for detailed *in vivo* imaging (especially in the *casper* strain), can be used for small molecule screens, and has a wide variety of cancer models available (Heilmann et al., 2015; Patton et al., 2021; Zhang et al., 2012). For melanoma, it has been widely used to study the effects of BRAF or NRAS (key initiating events in melanoma) and has uncovered important developmental and microenvironmental influences on these tumors (Kaufman et al., 2016; Zhang et al., 2018).

While less studied, the zebrafish has also been increasingly used to study host physiology and disease in the context of cancer. This includes metabolite tracing and inter-organ crosstalk (Naser et al., 2021). Several models of obesity have been generated in the zebrafish, using either diet or genetic manipulation, and closely resembles key aspects of the condition in humans (Landgraf et al., 2017; Song and Cone, 2007; Zang et al., 2018). For example, AgRP is a neuropeptide secreted in the hypothalamus as part of the central melanocortin signaling system that regulates food intake (Ollmann et al., 1997). Clinically, the central melanocortin system is the pathway with the most mutations observed in genetically obese patients (Loos and Yeo, 2022; Mendes de Oliveira et al., 2021). There is increased expression of AgRP in response to obesity in both mice and humans (Katsuki et al., 2001; Shutter et al., 1997). Ubiquitous overexpression of AgRP has been shown to increase appetite, resulting in increased weight and obesity in mice (Graham et al., 1997). The central melanocortin system and AgRP are highly conserved between mammals and zebrafish and overexpression of the peptide leads to a similar phenotype in this organism (Song and Cone, 2007; Song et al., 2003). The cells affected in these zebrafish models continue to be characterized, but at a minimum pertain to white adipocytes (i.e. subcutaneous or visceral adipocytes), since the fish are not thought to have thermogenic brown adipocytes. Other organs affected by obesity, including the liver and skeletal muscle, are also present in this organism, implying it can be readily studied in the context of cancer. In this study, we combine these unique attributes of the zebrafish to study the intersection of germline and somatic genetics in a melanoma model of obesity. The methods described here can be leveraged in future studies for larger scale discovery-based efforts to uncover new mediators of this crosstalk with relevance to the human diseases.

## Results

### AgRP overexpression promotes obesity in casper zebrafish

Previous studies have shown that overexpression of the orexigenic peptide Agouti-related protein AgRP1 in zebrafish results in fish that are overweight with hypertriglycerdemia and fatty liver (Song and Cone, 2007). We generated a plasmid in which the ubiquitin B (ubb) promoter drives the expression of zebrafish AgRP1 cDNA (zAgRP1) followed by a 2aGFP in order to visualize its expression (**Fig. 1A**) (Mosimann et al., 2011; Song et al., 2003). We first confirmed the obesogenic effects of zAgRP1 in fish without melanoma by injecting the plasmid into *casper* fish (*mpv17-/-, mitfa -/-; p53-/-*) (**Fig. 1B**). These fish also contain an inactive BRAF^V600E^ oncogene in the germline, so on their own will not develop melanoma but can be induced to so as explained further below. We found that in these non-melanoma fish, mosaic overexpression in this line led to an increase in weight in these fish over time, statistically observed at 4 months post fertilization (**Fig. 1C**). Furthermore, when observing male and female fish separately, we see that mosaic overexpression of zAgRP1 leads to increased weight in females at 4 months and a trend for an increase in weight in males (p=0.0585) at this time point (**Fig. 1D-E**). We outcrossed these fish to generate stable lines (*Tg(−3*.*5ubb:zAgRP1-2A-EGFP))*. In the F3 generation, both male and female zAgRP1 overexpressing fish have increased weight and are larger compared to wildtype quads (**Fig. 1F-H**). We did not observe differences in length, an alternative measure of obesity in zebrafish, in either the mosaic or stable *Tg(−3*.*5ubb:zAgRP1-2A-EGFP)* fish (**Fig. S1A-S1E**) (Song and Cone, 2007).

**Figure 1.**
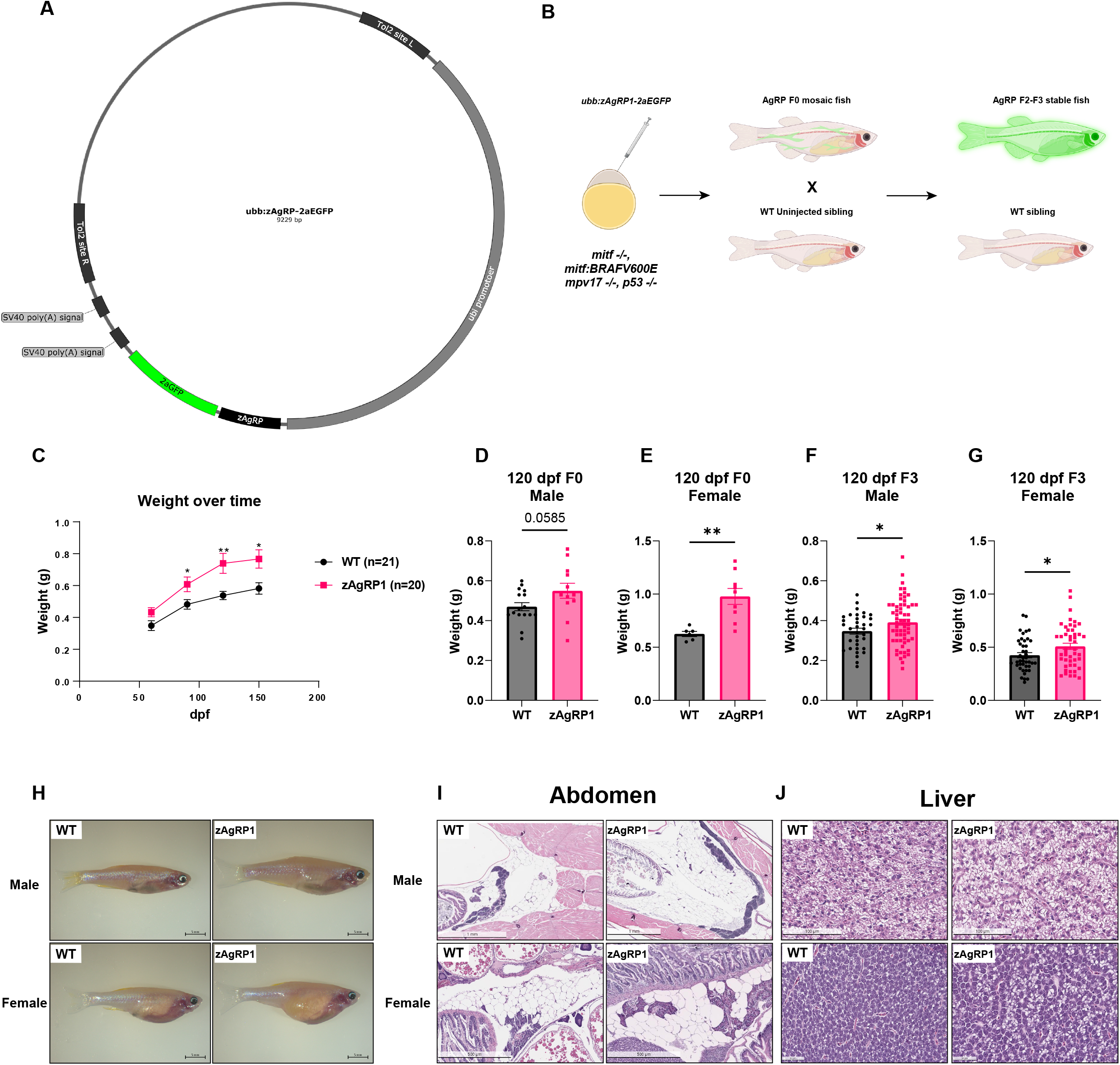
AgRP overexpression promotes obesity in casper triple zebrafish. A) AgRP overexpression construct driven by the ubb promoter. Created with Snapgene. (B) Schematic of generation of zAgRP1 overexpressing casper zebrafish (mpv17-/-, mitfa -/-; p53-/-, mitfa:BRAFV600E). Created with Biorender. (C) Weight of both male and female F0 fish combined over 6 months. (D-E) Weights of male (D) and female (E) at 120 dpf of F0 mosaic fish. Fish were separated into equal numbers at one month and weights measured at indicated time points. n=5-10 fish per genotype per biological replicate. Data is the average of 3 biological replicates. (F-G) Weights of male (F) and female (G) fish at 120 dpf of F3 stable line. (H) Representative images of male and female zAgRP1 fish and wild-type siblings. The data is the weight of all the fish from 2 separate clutches of fish across six separate tanks. n=35-60 fish per group. zAgRP1 fish and wild-type siblings were housed in the same tanks and identified via GFP fluorescence. (I-J) Histology of abdomen (I) and liver (J) from 7 month old F3 male and female zAgRP1 or wildtype siblings. Representative images. 2 fish per genotype per sex were sectioned for histololgy. * p ≤ 0.05, ** p ≤ 0.01 Welch’s t-test.

To further confirm that overexpression of zAgRP1 in these fish exhibited pathologic effects of obesity, we sectioned 5 month old F3 *Tg(−3*.*5ubb:zAgRP1-2A-EGFP)* fish and sent them for histology. We found that both male and female *Tg(−3*.*5ubb:zAgRP1-2A-EGFP)* fish had more abdominal adipose tissue compared to wildtype control fish (**Fig. 1I**). Furthermore, when observing the effect of zAgRP1 overexpression on the histology of the liver, we find that *Tg(−3*.*5ubb:zAgRP1-2A-EGFP)* fish exhibit fatty liver while the wildtype fish do not (**Fig. 1J**). Taken together this data demonstrate that overexpression of zAgRP1 driven by the ubiquitin promoter results in obesity in the *casper* strain of zebrafish, consistent with previous work overexpressing zAgRP1 driven under the actin promoter in zebrafish (Song and Cone, 2007).

### AgRP overexpression increases visceral adiposity

We wanted to further characterize the effect of zAgRP1 expression on overall adiposity and adipocyte dynamics in living fish. To do this, we first utilized the fluorescent dye BODIPY to visualize the fat depots in the fish in response to zAgRP1 overexpression. These dyes have been used extensively to study the anatomical distribution and amount of adipose tissue in living zebrafish (Minchin and Rawls, 2017). Upon staining both male and female zebrafish with BODIPY, we found that zAgRP1 overexpression increases overall adiposity compared to wildtype controls (**Fig. 2A-B**). Further analysis into the visceral and subcutaneous fat demonstrate that zAgRP1 specifically increases the area of adipose tissue in the visceral abdominal region and not the subcutaneous tail depot (**Fig. 2C-F**).This demonstrates that our *Tg(−3*.*5ubb:zAgRP1-2A-EGFP)* fish model, represents clinical aspects of obesity as the disease related adverse effects are more strongly associated with excess visceral fat (Fox et al., 2007; Goodpaster et al., 2003).

**Figure 2.**
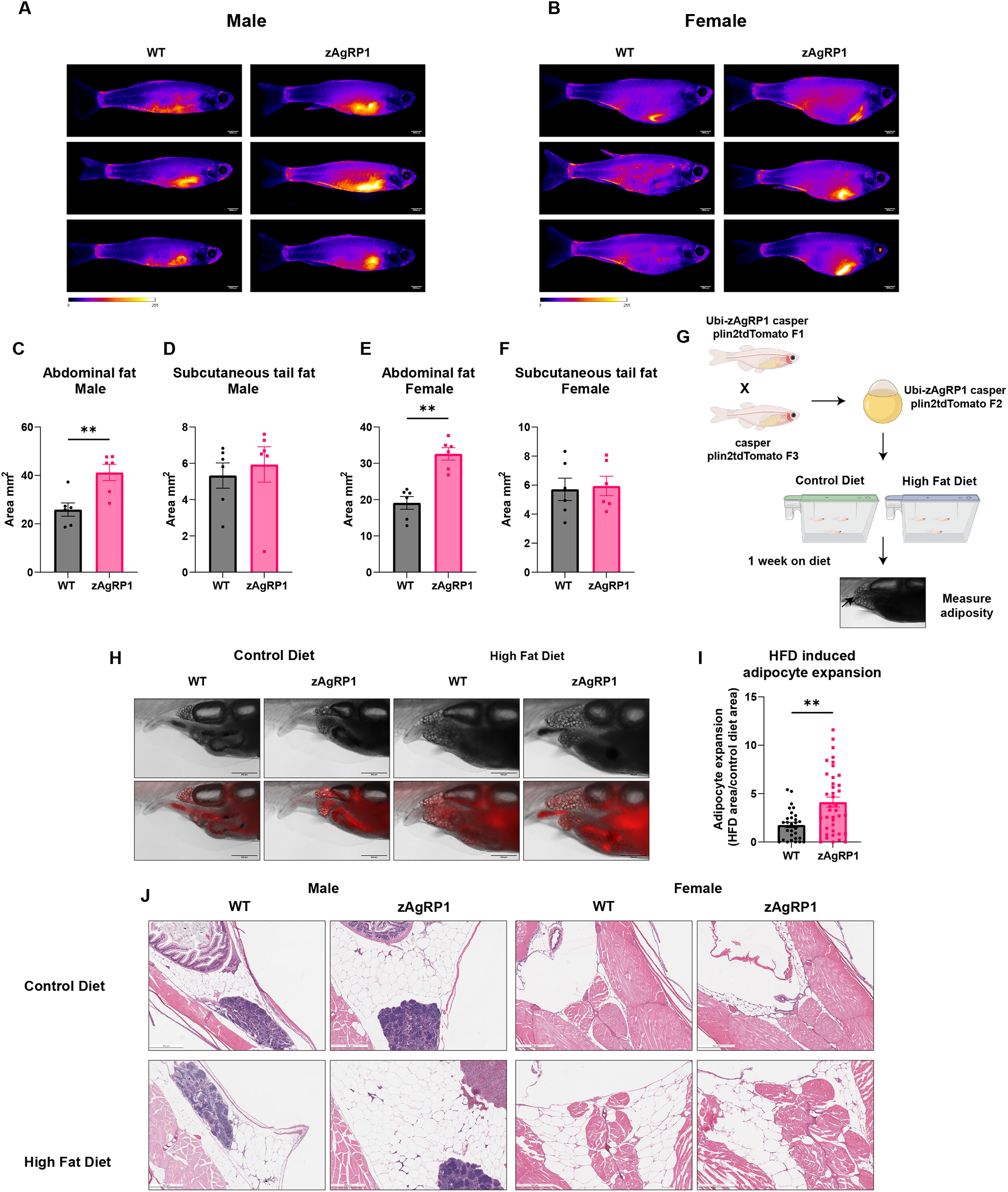
AgRP overexpression increases visceral adiposity and susceptibility to HFD. A-B) Representative images of male (A) and female (B) BODIPY stained fish. (C-F) Quantification of visceral abdominal (C, E) and subcutaneous (D, F) fat depots from BODIPY stained male (C, D) and female (E, F) fish. n=6 fish per condition over 3 biological replicates. (G) Schematic of plin2tdTomato HFD experiment. 21 dpf zAgRP1 or wild-type plin2tdTomato fish were put on either a control or high fat diet for one week and visceral adiposity measured. Created with Biorender. (H) Representative images of plin2tdTomato fish. (I) Quantification of adipocyte expansion. HFD induced adipocyte expansion was calculated by taking the ratio of the area of adipocyte tissue on control diet versus HFD for each genotype. n ≥ 30 fish per genotype across 3 biological replicates. (J) Histology of a cross section male and female adult quad fish with zAgRP1 or wild type controls on a high fat or control diet for 3 months. 2 fish per condition were sent for sectioning. ** p ≤ 0.01, Mann-Whitney test.

### AgRP results in increased susceptibility to HFD

Obesity has both a genetic and environmental component, and it is thought that patients with mutations in the melanocortin signaling pathway exhibit poor dietary control by preferring foods with a high fat content (van der Klaauw et al., 2016). Similarly studies in mice demonstrate that alterations in this pathway promote increased preference for high fat diets (HFD) (Koegler et al., 1999; Tung et al., 2007). Since AgRP plays a role in regulating adiposity as well as high fat diet seeking behaviors, we wanted to determine the effect of zAgRP1 overexpression in the zebrafish on adipocyte dynamics in the context of a HFD. To better visualize these dynamics (compared to BODIPY), we utilized a previously developed zebrafish line in which the adipocytes lipid droplets are fluorescently labeled with a plin2-tdTomato construct (Lumaquin et al., 2021) and are highly sensitive to HFD. We injected the *ubb:zAgRP1-2A-EGFP* construct into the *Tg(−3*.*5ubb:plin2-tdTomato)* line and outcrossed several generations to generate a stable line (**Fig. 2G**). We then took 21 dpf larvae and fed them either a commercially available HFD or control diet for a week and measured visceral adiposity using the tdTomato fluorescence (**Fig. 2H**). We found that in response to a HFD, *Tg(−3*.*5ubb:zAgRP1-2A-EGFP)* fish have increased adipocyte expansion compared to wildtype controls (**Fig. 2I**).

In order to confirm that this effect is conserved when fish are adults over a longer period of time, we fed the F3 *Tg(−3*.*5ubb:zAgRP1-2A-EGFP)* fish with a HFD for 3 months and then sent them for histology. We found that *Tg(−3*.*5ubb:zAgRP1-2A-EGFP)* fish that were fed a HFD had more adipose tissue as well as worse fatty liver compared to wildtype siblings on either diet alone or *Tg(−3*.*5ubb:zAgRP1-2A-EGFP)* fish on the control diet (**Fig. 2J and S2A**). Interestingly, we do not see a synergistic effect of HFD and zAgRP1 on adult fish weight over several weeks (**Fig. S3A-B**). Taken together, these findings demonstrate that zAgRP1 expressing fish are more susceptible to HFD induced obesity, further demonstrating the similarities between our model and the obesity phenotype observed in patients.

### AgRP increases melanoma onset in male zebrafish

Given the obesogenic effects of both AgRP and HFD, we next studied this in the context of melanoma. As described above, the transgenic zebrafish had a latent BRAF^V600E^ gene in the germline which can be activated by injection of the MiniCoopR plasmid, as previously described (Iyengar et al., 2012). We co-injected MiniCoopR-tdTomato with either the *ubb:zAgRP1-2A-EGFP* or an empty vector construct (**Fig. 3A**). Similar to the experiments above, the fish that were injected with *ubb:zAgRP1-2A-EGFP* are larger compared to empty vector controls (**Fig. 3B-C**). We screened the fish starting at 2 mpf, imaged the fish starting at 3 mpf, and assessed melanoma onset using a previously established rubric developed in the lab (Weiss et al., 2020). These criteria are based on the combination of hyperpigmentation, tdTomato fluorescence and growth of lesions into or out of the fish. This showed that the zAgRP1 overexpressing obese zebrafish develop tumors significantly faster than empty vector controls (**Fig. 3D**). When we segregated these fish by sex and calculated disease-free survival, we see that the increased onset is primarily seen in male fish but absent in females (**Fig. 3E-F**). These data are the first animal studies to demonstrate the clinically observed sex specific effect of obesity on melanoma (Sergentanis et al., 2013).

**Figure 3.**
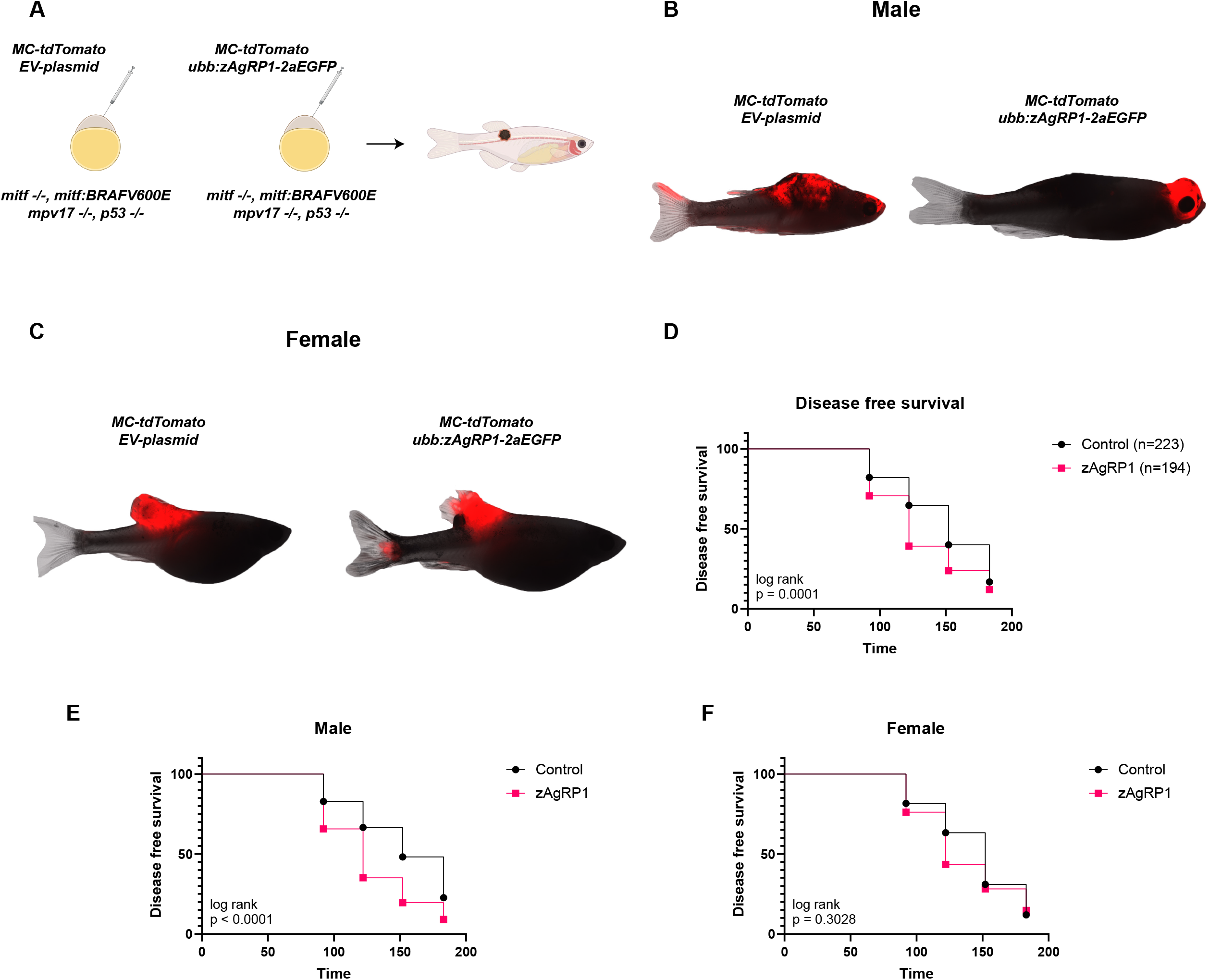
AgRP overexpression increases tumor onset in embryo injection model of melanoma. A) Schematic of embryo injection experiment to determine tumor onset with and without zAgRP1. Created with Biorender. (B-C) Representative images of male (B) and female (C) fish with tumors from EV control or zAgRP1 overexpression. (D) Disease free survival of MiniCoopR rescued male and female fish combined. (E-F) Disease free survival of MiniCoopR rescued male (E) and female (F) fish separated from D. Fish were injected at the one cell stage and monitored for tumors starting at 2 mpf at indicated time points. Data is the average of 3 biological replicates.

### AgRP increases tumor initiation and area in RB1 mutant melanoma

While this data clearly demonstrates that zAgRP1 overexpression leads to increased tumor onset, there are several limitations to this model in which the transgenes are injected at the embryonic stage. In this embryonic injection transgenic model, tumorigenesis occurs in the embryo and tumor onset manifests over time. This is an issue for this particular question, as zebrafish do not start differentiating their sex until ∼19 dpf and do not overtly display sex phenotypes until about one month post fertilization. Second, zAgRP1 overexpression induces overfeeding, which is not detected in weight until 4 mpf, long after the transgene is activated. Third, it is hard to detect differences in tumor size in this model, or study metastasis since the transgene is expressed everywhere and we cannot discern multifocal primary tumors versus metastatic lesions. To better address these issues, we instead turned to an electroporation based method called TEAZ (Transgene Electroporation in Adult Zebrafish (Callahan et al., 2018)), in which plasmids are directly electroporated into adult skin cells. Thus, tumor onset occurs somatically in adulthood (similar to humans), is spatially and temporally controlled, and can be monitored for tumor area and metastasis occurrence.

We used TEAZ to initiate tumors driven by BRAFV600E;p53-/-;RB1-/-with or without zAgRP1 overexpression (**Fig. 4A**). This approach had been previously shown to induce melanoma development and is responsive to genetic perturbations (Baggiolini et al., 2021; Tagore et al., 2021). We found that zAgRP1 increases tumor initiation detected at 14 dpe (days post-electroporation) as well as tumor area of early (14 dpe **Fig. 4B-C**) and late lesions (42 dpe, **Fig. 4D-E**). We have found that these lesions are relatively slow growing and rarely develop metastasis. Therefore, we sought to determine the effect of zAgRP1 on a more aggressive model by swapping the RB1 deletion for a PTEN deletion (**Fig. S4A**). We have found that these tumors are more aggressive, growing faster and larger compared to the RB1 deletion, a phenomenon that has previously been reported in mice (Dankort et al., 2009). Interestingly when we initiate tumors using TEAZ with a PTEN deletion, we find that indeed these tumors are larger and grow faster than their RB1 counterparts, however there was no significant effect of zAgRP1 layered on top of a PTEN deletion (**Fig. S4B-E**). This was the case for both early and late lesions.

**Figure 4.**
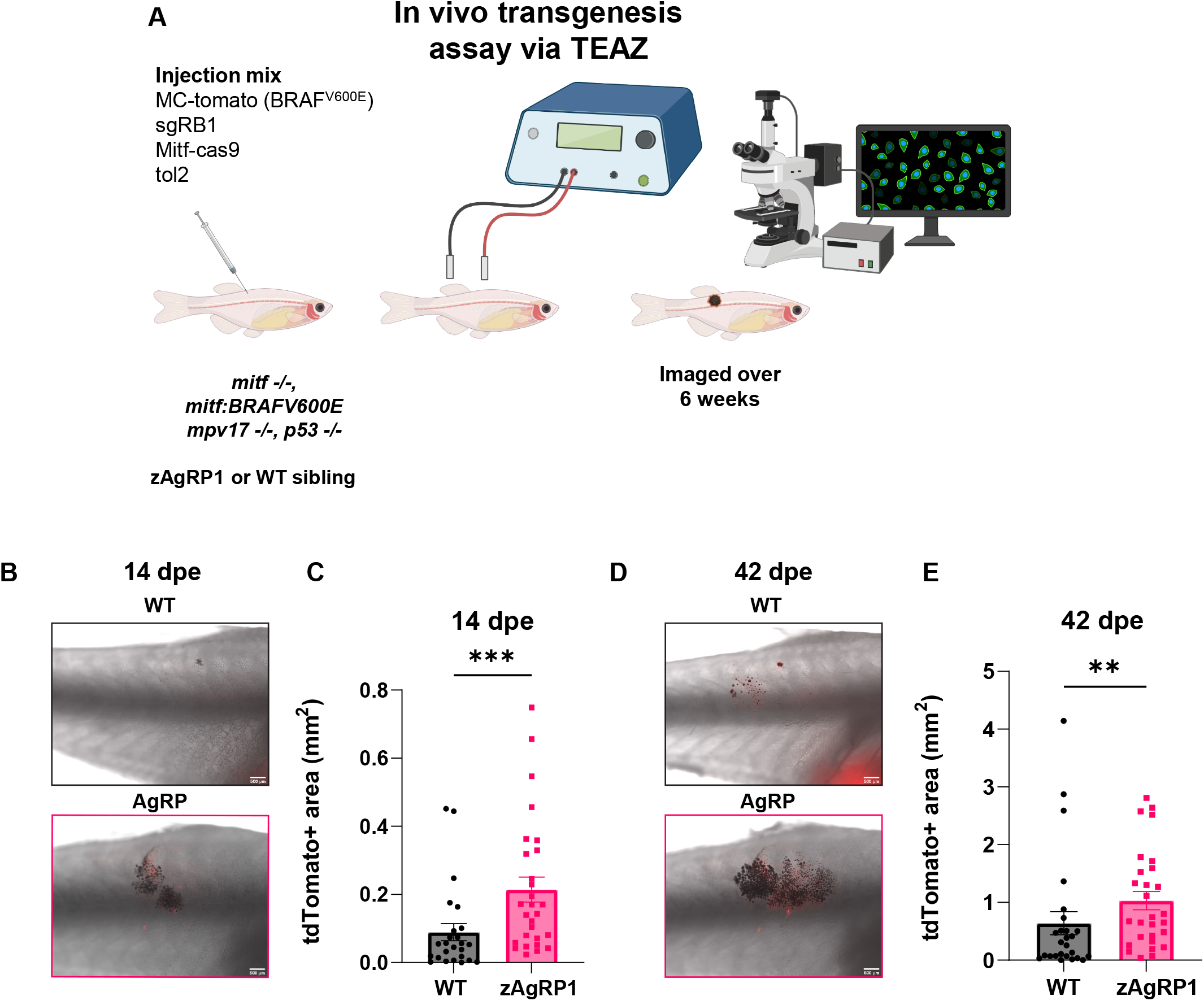
Obesity increases tumor initiation and area in electroporation model of RB1 mutant melanoma. A) Schematic of in vivo transgenesis assay via TEAZ. Adult casper (mitfa:BRAFV600E, p53-/-, mitfa-/-, mpv17-/-) zAgRP1 or WT F0 fish were injected with MiniCoopR-tdTomato, sgRB1, mitfa:Cas9, and tol2 constructs and then electroporated. Fish were analyzed for tumor initiation and area by fluorescence microscopy over 6 weeks. Created with Biorender. (B-C) Tumor area at 14 dpe. Representative images (B) and quantification (C) of WT and zAgRP1 fish. (D-E) Tumor area at 42 dpe. Representative images (D) and quantification (E) of WT and zAgRP1 fish. n ≥ 25 per genotype. Data is the average of 3 biological replicates. ** p ≤ 0.01, *** p ≤ 0.001, Mann-Whitney test.

### Obesity increases tumor initiation in RB1 mutant melanoma in male zebrafish

Since the TEAZ transgenesis assay allows us to induce a tumor in a fully immunocompetent adult animal, we next sought to extend our findings on obesity and melanoma in the context of zAgRP1 overexpression or high fat diet and whether this was dependent on sex. We used the *Tg(−3*.*5ubb:zAgRP1-2A-EGFP)* WT siblings and fed them a HFD for one week based on our data that zAgRP1 can induce changes in 1 mpf larvae after a week on the diet (**Fig. 2H-I**). We then introduced the BRAF^V600E^ mutation in conjunction with a RB1 deletion into the fish using TEAZ and monitored fish for tumors via fluorescence microscopy over the course of 12 weeks (**Fig. 5A**). During this time the fish remained on their respective diets. This revealed a significant sex-based difference in tumor growth. In male fish, all obesity conditions (genetic, diet or combined) led to tumors that were larger compared to WT control diet fish. When observing early (21 dpe, **Fig. 5D and H**) and late (63 dpe, **Fig. 5E and H**) individual time points, we see that the effect is stronger earlier on and seems to become less apparent at later stages, suggesting that the effect is primarily on tumor initiation more so than progression. In contrast, in females we did not observe a significant difference in growth curves between the WT control fish and either zAgRP1 or HFD alone at either early (21 dpe, **Fig. 5F and H**) or late (63 dpe, **Fig. 5G and H**) time points. However, the combination of genetic and diet induced obesity did have an additive effect on tumor area in combination only at early time points (21 days), but this is not seen in later lesions, further underscoring lack of increased tumor area in response to obesity specifically in females.

**Figure 5.**
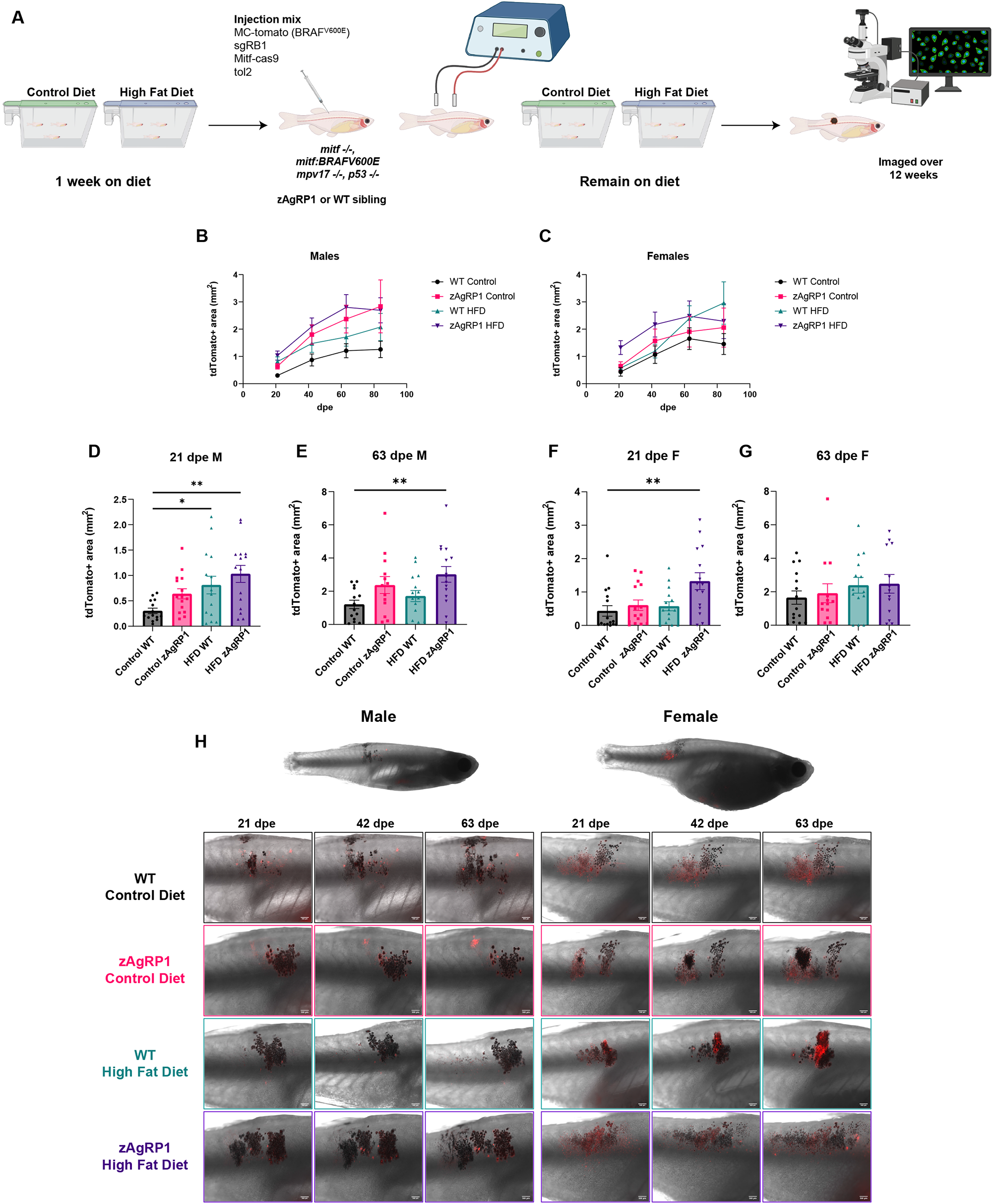
Obesity increases tumor growth in sex dependent manner in RB1 mutant melanoma. (A) Schematic of in vivo transgenesis assay via TEAZ with the addition of a high fat diet. Fish were put on a control or high fat diet for one week. Adult casper (mitfa:BRAFV600E, p53-/-, mitfa-/-, mpv17-/-) zAgRP1 or WT F3 fish were injected with MiniCoopR-tdTomato, sgRB1, mitfa:Cas9, and tol2 constructs and then electroporated. Fish were analyzed for tumor initiation and area by fluorescence microscopy over 12 weeks while they remained on their respective diets. Created with Biorender. (B-C) Tumor growth as measured by tdTomato+ area over time. TdTomato+ area of lesions on zAgRP1 or WT male (B) and female (C) fish at indicated dpe. Growth curves were analyzed via Mixed-effects analysis, with p=0.0344 significance when comparing the different conditions in males and p=0.2391 in females. (D-E) tdTomato+ area for early (21 dpe, D) and late (63 dpe, E) lesions in male zAgRP1 or WT fish on either a control or high fat diet. (F-G) tdTomato+ area for early (21 dpe, F) and late (63 dpe, G) lesions in female zAgRP1 or WT fish on either a control or high fat diet. (H) Representative fluorescence overlaid on brightfield images of lesions in male and female fish at indicated time dpe. n=14 fish per condition per sex and is the average of 3 biological replicates. * p ≤ 0.05, ** p ≤ 0.01, Dunnet’s multiple comparisons test.

### Obesity-related effects on melanoma are not seen with PTEN

Finally, we determined whether this was maintained with a different genetic driver. We therefore studied the effect of BRAFV600E;p53-/-;PTEN-/-. Unlike the case for RB1-/-, we did not see a significant difference in the tumor growth in the context of a PTEN deletion in both male and female fish (**Fig. S5B-C**). This was true in both early and late lesions (**Fig. S5D-H**). As expected, the PTEN tumors were larger and grew faster compared to those with an RB1 deletion. Overall, these data demonstrate that the interaction between systemic alterations (i.e. diet, sex) and the tumor is dependent on the genotype of the tumor itself (PTEN vs RB1 deletion). This highlights that it is critical to develop models in which it is possible to study multiple facets in both the host and the tumor concurrently. The flexibility of the zebrafish system will allow future studies to dissect these interactions in a high-throughput manner.

## Discussion

In this study, we aimed to develop the zebrafish as a new model for studying the interaction between obesity and melanoma, and whether this depends on somatic and germline factors. This revealed a sex-specific effect on melanoma initiation that is consistent with clinical literature on human patients. This effect was most clear in tumors initiated by BRAFV600E;p53-/-;RB1-/-but not BRAFV600E-/-;p53-/-;PTEN-/-.

While studies such as these are more traditionally done using mouse models of cancer, the rapidity by which we can generate these perturbations, and precisely quantify tumor initiation and progression, make the zebrafish an ideal system for future studies. The major advance offered in this system is the ease by which we can make somatic mutations (i.e. loss of RB1 or PTEN) using the TEAZ system, which is far more rapid and simpler than comparable mouse models. Moreover, these can be readily combined with germline alterations in genes such as AgRP, one of the most frequent alterations seen in patients with obesity, or with dietary changes such as high-fat diets. One of the important observations in our research is that obesity has a sex-specific effect on melanoma initiation, with obese males having a greater effect than obese females. However, despite numerous human and animal studies demonstrating that males are at higher risk of developing melanoma (Pandey et al., 2012; Renehan et al., 2008; Sergentanis et al., 2013), it has also long been observed that females have improved survival once they have the disease. The mechanisms regulating this discrepancy remain poorly understood. Potential reasons include a relatively stronger immune responsiveness in females (Bouman et al., 2005; Gubbels Bupp et al., 2018). Hormonal influences are also thought to play an important, yet still not understood role. Estrogen binds to multiple receptors with somewhat opposing effects. Estrogen receptor α (ERα) is thought to be pro-proliferative, whereas Estrogen receptor β (ERβ) may act in an anti-proliferative manner and men were found to have comparatively lower levels of ERβ (de Giorgi et al., 2013). Estrogens can also bind to the more recently described G protein-coupled estrogen receptor (GPER), which can inhibit melanoma growth and are thought to mediate some of the protective effect in females (Natale et al., 2018). Testosterone likely also plays a role, since melanomas are known to express androgen receptors (AR), with dihydrotestosterone thought to be especially mitogenic (Richardson et al., 1999). Adding to this complexity is the so-called “obesity paradox”, in which obesity can be associated with improved survival specifically in males. For example, a large retrospective meta-analysis showed that obese males have improved response to targeted or immunotherapy treatments (McQuade et al., 2018). While our studies do not address the mechanisms involved in these seemingly contradictory effects on initiation versus survival, there is evidence that polymorphisms of p53 may play a role in such sex-specific effects (Oliveira et al., 2017). The fact that we see this sex difference in the context of p53/RB1, but not p53/PTEN, also highlights a potential interaction between germline alleles and somatic mutations in the PTEN/PI3K pathway. This is particularly compelling given the known interactions between ERβ and the PI3K pathway (Dey et al., 2013; Lei et al., 2020). These would be fruitful areas for future exploration using the zebrafish system we have developed.

## Materials and Methods

### Zebrafish husbandry

All zebrafish (*Danio rerio*) were bred and housed in the Aquatics facility in Zuckerman Research Center at Sloan Kettering Institute. All fish were housed in standard conditions with a water temperature of 28.5°C, controlled salinity, a pH of 7.4, and a light/dark cycle of 14hrs/10hrs. Unless otherwise indicated, fish were initially fed rotifers followed by a standard commercial pellet diet (GEMMA) 3 times per day. Fish were housed at a density no higher than 10 fish per liter. Anesthesia of adult zebrafish was carried out using Tricaine (4g/L, Syndel USA) that was diluted to 0.16 mg/mL. All of the protocols listed in this manuscript were reviewed and approved by the Memorial Sloan Kettering Cancer Center Institutional Animal Care and Use Committee, protocol #12-05-008.

### Zebrafish transgenic lines

The zebrafish lines used in these studies were the casper (*mitfa:*BRAF^V600E^, *p53-/-, mitfa-/-, mpv17-/-)* fish *(Baggiolini et al., 2021*; Tagore et al., *2021*) and the *Tg(−3*.*5ubb:plin2-tdTomato)* line (Lumaquin et al., 2021). These lines were generated in the white lab previously. For this manuscript we also generated the *Tg(−3*.*5ubb:zAgRP1-2A-EGFP)* casper (*mitfa:*BRAF^V600E^, *p53-/-, mitfa-/-, mpv17-/-)* fish and the *Tg(−3*.*5ubb:plin2-tdTomato, -3*.*5ubb:zAgRP1-2A-EGFP)* casper fish lines as described below.

### Generation of AgRP Construct

To generate the *ubb:zAgRP1-2A-EGFP* construct, Gateway Cloning, using the LR Gateway Enzyme mix (Thermo Fisher, Waltham, USA; catalog #11791019), was completed using a p5E-ubb, pME-zAgRP1, p3E-2A-EGFP into the pDestTol2pA2-blastocidin destination vector. The control plasmid, an empty pDestTol2pA2-blastocidin vector, was generated from colonies that arose in the destination vector control reaction. All selected colonies were cultured overnight and prepared using the HiSpeed Plasmid Maxi Kit (Qiagen, Hilden Germany, cat# 12663) and sequenced at Genewiz (South Plainfield NJ USA) for confirmation. To create the pME-zAgRP1 construct for gateway cloning, the cDNA sequence was obtained from (Song et al., 2003) and ordered as a gBlock Gene Fragment (IDT DNA, Coralville IA USA), PCR amplified using AccuPrime Taq DNA Polymerase High Fidelity (Invitrogen, Waltham MA USA, cat# 12346086), and gel extracted using the NucleoSpin Gel and PCR Clean-up kit (Takara Bio USA Inc, San Jose CA USA, cat# 740609). The fragment was subsequently cloned into the pME vector using the pENTR/D-TOPO Cloning kit (Invitrogen, Waltham MA USA, cat# K240020). Colonies were prepared using the Qiaprep Spin Miniprep kit (QiAgen, Hilden Germany, cat# 27106) and sequenced at Genewiz (South Plainfield NJ USA) for confirmation.

### Generation of AgRP fish

In order to generate the *Tg(−3*.*5ubb:zAgRP1-2A-EGFP)* casper (*mitfa:*BRAF^V600E^, *p53-/-, mitfa-/-, mpv17-/-)* mosaic fish, the *ubb:zAgRP1-2A-EGFP* and Tol2 mRNA were injected into the yolk of casper (*mitf:*BRAF^V600E^, *p53-/-, mitf-/-, mpv17-/-)* fish at the one cell stage (Baggiolini et al., 2021; Tagore et al., 2021). Fish were sorted for GFP+ fluorescence due to the 2aEGFP and used for mosaic experiments or to generate a stable line of F2 and F3 fish. For mosaic experiments, uninjected quad fish from the same clutch were used as control. To make the stable line, F0 fish were out crossed 2 generations to quad zebrafish, and GFP-siblings were used as the WT control.

### Generation of AgRP Plin2tdTomato fish

In order to generate the *Tg(−3*.*5ubb:plin2-tdTomato, -3*.*5ubb:zAgRP1-2A-EGFP)* casper mosaic fish, 25 ng/μL of the *ubb:zAgRP1-2A-EGFP* and 20 ng/μL of Tol2 mRNA were injected into the yolk of *Tg(−3*.*5ubb:plin2-tdTomato)* casper fish at the one cell stage (Lumaquin et al., 2021). Fish were sorted for both overall GFP+ fluorescence and GFP+ hearts used to generate a stable line of F2 and F3 fish. To make the stable line, F0 fish were out-crossed 2 generations to *Tg(−3*.*5ubb:plin2-tdTomato)* casper zebrafish, and GFP-body, but GFP+ heart siblings were used as the WT control.

### Determination of zebrafish weight and length

Adult fish were anesthetized with tricaine and then dried using a paper towel. Length was determined to the closest 0.01 mm using electronic calipers (Cole-Parmer, Vernon Hills IL USA, Cat# # EW-09925-43) and weight to the nearest 0.01 g using a portable balance (OHAUS, Parsippany NJ USA, Cat# SPX222). Fish were then placed into a tank with fresh system water to recover.

### Histology

Zebrafish were sacrificed using ice-cold water. Head and tail were removed, and fish were placed in 4% PFA in PBS for 72 hours at 4°C on a rocker. Fish were then transferred to 70% EtOH for 24 hours at 4°C on a rocker. Fish were sent to Histowiz (Brooklyn NY USA), where they were paraffin embedded, sectioned coronally at 5 μm, and underwent hematoxylin and eosin staining.

### High Fat Diet Feeding

For high fat diet experiments, fish were fed a commercially developed pelleted fish food (Sparos, Portugal) (Lumaquin et al., 2021). Both the HFD and matched control diets were developed by Sparos. The crude composition as per fed basis for the control diet was: 57.3% crude protein, 13.1% crude fat, 0.5 % fiber, 8.8% ash, and 20.3 MJ/kg gross energy. The crude composition as per fed basis for the HFD was: 57.3% crude protein, 24.8% crude fat, 0.5 % fiber, 8.8% ash, and 23.3 MJ/kg gross energy. The diets were analyzed for final composition. The Sparos control diet contains 30% fishmeal, 33% squid meal, 5% fish gelatin, 5.5% wheat gluten, 12% cellulose, 2.5% soybean oil, 2.5% rapeseed oil, 2% vitamins and minerals, 0.1% vitamin E, 0.4% antioxidant, 2% monocalcium phosphate, and 2.2% calcium silicate. The Sparos HFD contains 30% fishmeal, 33% squid meal, 5% fish gelatin, 5.5% wheat gluten, 12% palm oil, 2.5% soybean oil, 2.5% rapeseed oil, 2% vitamins and minerals, 0.1% vitamin E, 0.4% antioxidant, 2% monocalcium phosphate, and 2.2% calcium silicate. For the larvae experiments, 21 dpf larvae were moved to 0.8L tanks in equal density (15-20 larvae). Fish were fed 0.1g of food split between 2x per day. Fish were kept on the diet for 1 week and then imaged for visceral adiposity via tdTomato expression. The images were analyzed and the visceral adiposity measured using MATLAB (Lumaquin et al., 2021). HFD induced expansion was calculated as the tdTomato+ area of the HFD fed group divided by the average of the control fed group for the indicated genotype. For adult fish experiments, fish were fed 5% of bodyweight per fish split over 2x per day. Fish were kept on the diet for indicated time points (1-3 months).

### Embryo injection transgenesis and analysis

In order to generate melanomas using embryo injection based transgenics, we used the MiniCoopR based transgenic model as previously described (Ceol et al., 2011; Kaufman et al., 2016). Fish were injected at the one-cell stage with 15 ng/μL of MiniCoopR-tdTomato, 15 ng/μL of either *ubb:zAgRP1-2A-EGFP* or empty vector control, and 20 ng/μL Tol2 mRNA. Fish were monitored at 48 hrs post fertilization for GFP and tdTomato fluorescence. Fish were put in the nursery at 5 dpf and then monitoring for tumors starting at 2 mpf, and imaging at 3 mpf. Fish were checked monthly for tumors up until 6 mpf. Tumors were determined based on a rubric of criteria developed in the lab previously (Weiss et al., 2020). The basis of the criteria is hyperpigmentation, tdTomato fluorescence and growth into or out of the fish. Disease Free Survival curves were generated using GraphPad Prism 8 (Graphpad, San Diego, USA).

### TEAZ based transgenesis and analysis

Melanomas were generated using TEAZ as described previously (Callahan et al., 2018). All tumors generated had a *BRAF*^*V600E*^ mutation and loss of *p53* with the additional loss of either *rb1* or *ptena* and *ptenb*. To generate these tumors, fish were anesthetized with 0.16 mg/mL tricaine and injected with 1uL of rb1 tumor plasmid mix (250 ng/μL *mitf:*Cas9, 250 ng/μL MiniCoopR-tdTomato, 106 ng/μL sgRB1, 67 ng/μL Tol2) or ptena/b tumor plasmid mix (250 ng/μL *mitf:*Cas9, 250 ng/μL MiniCoopR-tdTomato, 27.3 ng/μL gPTENa, 27.3 ng/μL gPTENb, 61 ng/μL Tol2) into the skin below the dorsal fin. Fish were then electroporated and returned to fresh system water to recover. Electroporation was carried out using the CM 830 Electro Square Porator (BTX Harvard Apparatus, Holliston MA USA) and the 3mm platinum Tweezertrodes (BTX Harvard Apparatus, Holliston MA USA, Cat# 45-0487). The electroporator settings were LV mode with a voltage of 40 V, 5 pulses, 60 ms pulse length and 1 s pulse interval. Electroporated zebrafish were imaged serially for up to 12 weeks dpe using brightfield and fluorescence imaging. Area of tdTomato fluorescence was quantified at indicated time points using FIJI.

### Imaging and Analysis

Zebrafish were imaged using an upright Zeiss AxioZoom V16 Fluorescence Stereo Zoom Microscope equipped with a motorized stage, brightfield and fluorescent filter sets (mCherry or Cy5, GFP, and tdTomato). Adult fish were imaged with a x0.5 adjustable objective lens and larvae were imaged with a x1.0 adjustable objective lens. To acquire images, zebrafish were lightly anesthetized with 0.16 mg/mL tricaine. Images were acquired with the Zeiss Zen Pro v2 and exported as CZI files for visualization and analysis. Imaging analysis was completed manually using FIJI or automated with MATLAB software (Mathworks, Natick, USA).

### Statistical Analysis

All experiments were completed at least three times as independent biological experiments. Statistical analysis was completed using Graphpad Prism versions 8 and 9 (Graphpad, San Diego, USA). Exact sample sizes and statistical tests for each experiment are detailed in the figure legends. Data are represented as mean ± standard error of mean (SEM). Statistical significance was set at p≤0.05.

## Acknowledgements

We thank members at the Memorial Sloan Kettering Cancer Center Aquatics Core. We would also like to thank the rest of the White Lab members for their thoughtful insights on the project during many lab meetings.

## Competing Interests

EM, DL, YM, and SS have no competing interests.

RMW Is a paid consultant to N-of-One Therapeutics, a subsidiary of Qiagen. Receives royalty payments for the use of the casper zebrafish line from Carolina Biologicals.

## Funding

This work was funded by the National Institute of Health Individual Predoctoral to Postdoctoral Fellow Transition Award 5K00CA223016-04 (to E.M.), the Kirschstein-NRSA predoctoral fellowship F30CA254152 (to D.L.), the Kirschstein-NRSA predoctoral fellowship F30CA265124 (to Y.M.), the Melanoma Research Foundation Career Development Award grant number 719502 and an MSKCC TROT T32 training grant T32 CA16000 (to S.S), the Medical Scientist Training Program T32GM007739-42 (to D.L. and Y.M.), This research was funded in part through the NIH/NCI Cancer Center Support Grant P30 CA008748; Melanoma Research Alliance, The Debra and Leon Black Family Foundation, NIH Research Program Grants R01CA229215 and R01CA238317, NIH Director’s New Innovator Award DP2CA186572, The Pershing Square Sohn Foundation, The Mark Foundation for Cancer Research, The American Cancer Society (all to R.M.W.).

## Author Contributions

EM and RMW developed the experiments, interpreted the results, and wrote the manuscript. EM performed all experiments, data collection and analysis. DL helped with development and analysis of plin2 reporter fish. EM, YM, and SS helped with the development and analysis of the TEAZ melanoma model. RMW provided funding for the project and oversaw all experiments. All the authors read and edited the manuscript.

## References

Baggiolini, A., Callahan, S. J., Montal, E., Weiss, J. M., Trieu, T., Tagore, M. M., Tischfield, S. E., Walsh, R. M., Suresh, S., Fan, Y., et al. (2021). Developmental chromatin programs determine oncogenic competence in melanoma. Science 373, eabc1048.

Björntorp, P. and Sjöström, L. (1971). Number and size of adipose tissue fat cells in relation to metabolism in human obesity. Metabolism 20, 703–713.

Bouman, A., Heineman, M. J. and Faas, M. M. (2005). Sex hormones and the immune response in humans. Hum. Reprod. Update 11, 411–423.

Brandon, E. L., Gu, J.-W., Cantwell, L., He, Z., Wallace, G. and Hall, J. E. (2009). Obesity promotes melanoma tumor growth: role of leptin. Cancer Biol. Ther. 8, 1871–1879.

Callahan, S. J., Tepan, S., Zhang, Y. M., Lindsay, H., Burger, A., Campbell, N. R., Kim, I. S., Hollmann, T. J., Studer, L., Mosimann, C., et al. (2018). Cancer modeling by Transgene Electroporation in Adult Zebrafish (TEAZ). Dis. Model. Mech. 11,.

Ceol, C. J., Houvras, Y., Jane-Valbuena, J., Bilodeau, S., Orlando, D. A., Battisti, V., Fritsch, L., Lin, W. M., Hollmann, T. J., Ferré, F., et al. (2011). The histone methyltransferase SETDB1 is recurrently amplified in melanoma and accelerates its onset. Nature 471, 513– 517.

Dankort, D., Curley, D. P., Cartlidge, R. A., Nelson, B., Karnezis, A. N., Damsky, W. E., You, M. J., DePinho, R. A., McMahon, M. and Bosenberg, M. (2009). Braf(V600E) cooperates with Pten loss to induce metastatic melanoma. Nat. Genet. 41, 544–552.

Dey, P., Barros, R. P. A., Warner, M., Ström, A. and Gustafsson, J.-Å. (2013). Insight into the mechanisms of action of estrogen receptor β in the breast, prostate, colon, and CNS. J. Mol. Endocrinol. 51, T61–74.

de Giorgi, V., Gori, A., Gandini, S., Papi, F., Grazzini, M., Rossari, S., Simoni, A., Maio, V. and Massi, D. (2013). Oestrogen receptor beta and melanoma: a comparative study. Br. J. Dermatol. 168, 513–519.

Fox, C. S., Massaro, J. M., Hoffmann, U., Pou, K. M., Maurovich-Horvat, P., Liu, C.-Y., Vasan, R. S., Murabito, J. M., Meigs, J. B., Cupples, L. A., et al. (2007). Abdominal visceral and subcutaneous adipose tissue compartments: association with metabolic risk factors in the Framingham Heart Study. Circulation 116, 39–48.

Goodpaster, B. H., Krishnaswami, S., Resnick, H., Kelley, D. E., Haggerty, C., Harris, T. B., Schwartz, A. V., Kritchevsky, S. and Newman, A. B. (2003). Association between regional adipose tissue distribution and both type 2 diabetes and impaired glucose tolerance in elderly men and women. Diabetes Care 26, 372–379.

Graham, M., Shutter, J. R., Sarmiento, U., Sarosi, I. and Stark, K. L. (1997). Overexpression of Agrt leads to obesity in transgenic mice. Nat. Genet. 17, 273–274.

Gubbels Bupp, M. R., Potluri, T., Fink, A. L. and Klein, S. L. (2018). The confluence of sex hormones and aging on immunity. Front. Immunol. 9, 1269.

Hales, C. M., Carroll, M. D., Fryar, C. D. and Ogden, C. L. (2020). Prevalence of Obesity and Severe Obesity Among Adults: United States, 2017-2018. NCHS Data Brief 1–8.

Heilmann, S., Ratnakumar, K., Langdon, E., Kansler, E., Kim, I., Campbell, N. R., Perry, E., McMahon, A., Kaufman, C., van Rooijen, E., et al. (2015). A quantitative system for studying metastasis using transparent zebrafish. Cancer Res. 75, 4272–4282.

Iyengar, S., Houvras, Y. and Ceol, C. J. (2012). Screening for melanoma modifiers using a zebrafish autochthonous tumor model. J. Vis. Exp. e50086.

Karimi, K., Lindgren, T. H., Koch, C. A. and Brodell, R. T. (2016). Obesity as a risk factor for malignant melanoma and non-melanoma skin cancer. Rev. Endocr. Metab. Disord. 17, 389– 403.

Katsuki, A., Sumida, Y., Gabazza, E. C., Murashima, S., Tanaka, T., Furuta, M., Araki-Sasaki, R., Hori, Y., Nakatani, K., Yano, Y., et al. (2001). Plasma levels of agouti-related protein are increased in obese men. J. Clin. Endocrinol. Metab. 86, 1921–1924.

Kaufman, C. K., Mosimann, C., Fan, Z. P., Yang, S., Thomas, A. J., Ablain, J., Tan, J. L., Fogley, R. D., van Rooijen, E., Hagedorn, E. J., et al. (2016). A zebrafish melanoma model reveals emergence of neural crest identity during melanoma initiation. Science 351, aad2197.

Koegler, F. H., Schaffhauser, A. O., Mynatt, R. L., York, D. A. and Bray, G. A. (1999). Macronutrient diet intake of the lethal yellow agouti (Ay/a) mouse. Physiol. Behav. 67, 809– 812.

Landgraf, K., Schuster, S., Meusel, A., Garten, A., Riemer, T., Schleinitz, D., Kiess, W. and Körner, A. (2017). Short-term overfeeding of zebrafish with normal or high-fat diet as a model for the development of metabolically healthy versus unhealthy obesity. BMC Physiol. 17, 4.

Lei, S., Fan, P., Wang, M., Zhang, C., Jiang, Y., Huang, S., Fang, M., He, Z. and Wu, A. (2020). Elevated estrogen receptor β expression in triple negative breast cancer cells is associated with sensitivity to doxorubicin by inhibiting the PI3K/AKT/mTOR signaling pathway. Exp. Ther. Med. 20, 1630–1636.

Loos, R. J. F. and Yeo, G. S. H. (2022). The genetics of obesity: from discovery to biology. Nat. Rev. Genet. 23, 120–133.

Lumaquin, D., Johns, E., Montal, E., Weiss, J. M., Ola, D., Abuhashem, A. and White, R. M. (2021). An in vivo reporter for tracking lipid droplet dynamics in transparent zebrafish. eLife 10,.

McQuade, J. L., Daniel, C. R., Hess, K. R., Mak, C., Wang, D. Y., Rai, R. R., Park, J. J., Haydu, L. E., Spencer, C., Wongchenko, M., et al. (2018). Association of body-mass index and outcomes in patients with metastatic melanoma treated with targeted therapy, immunotherapy, or chemotherapy: a retrospective, multicohort analysis. Lancet Oncol. 19, 310–322.

Mendes de Oliveira, E., Keogh, J. M., Talbot, F., Henning, E., Ahmed, R., Perdikari, A., Bounds, R., Wasiluk, N., Ayinampudi, V., Barroso, I., et al. (2021). Obesity-Associated GNAS Mutations and the Melanocortin Pathway. N. Engl. J. Med. 385, 1581–1592.

Minchin, J. E. N. and Rawls, J. F. (2017). A classification system for zebrafish adipose tissues. Dis. Model. Mech. 10, 797–809.

Mosimann, C., Kaufman, C. K., Li, P., Pugach, E. K., Tamplin, O. J. and Zon, L. I. (2011). Ubiquitous transgene expression and Cre-based recombination driven by the ubiquitin promoter in zebrafish. Development 138, 169–177.

Naser, F. J., Jackstadt, M. M., Fowle-Grider, R., Spalding, J. L., Cho, K., Stancliffe, E., Doonan, S. R., Kramer, E. T., Yao, L., Krasnick, B., et al. (2021). Isotope tracing in adult zebrafish reveals alanine cycling between melanoma and liver. Cell Metab. 33, 1493–1504 e5.

Natale, C. A., Li, J., Zhang, J., Dahal, A., Dentchev, T., Stanger, B. Z. and Ridky, T. W. (2018). Activation of G protein-coupled estrogen receptor signaling inhibits melanoma and improves response to immune checkpoint blockade. eLife 7,.

Oliveira, C., Lourenço, G. J., Rinck-Junior, J. A., de Moraes, A. M. and Lima, C. S. P. (2017). Polymorphisms in apoptosis-related genes in cutaneous melanoma prognosis: sex disparity. Med. Oncol. 34, 19.

Ollmann, M. M., Wilson, B. D., Yang, Y. K., Kerns, J. A., Chen, Y., Gantz, I. and Barsh, G. S. (1997). Antagonism of central melanocortin receptors in vitro and in vivo by agouti-related protein. Science 278, 135–138.

Pandey, V., Vijayakumar, M. V., Ajay, A. K., Malvi, P. and Bhat, M. K. (2012). Diet-induced obesity increases melanoma progression: involvement of Cav-1 and FASN. Int. J. Cancer 130, 497–508.

Patton, E. E., Zon, L. I. and Langenau, D. M. (2021). Zebrafish disease models in drug discovery: from preclinical modelling to clinical trials. Nat. Rev. Drug Discov. 20, 611–628.

Renehan, A. G., Tyson, M., Egger, M., Heller, R. F. and Zwahlen, M. (2008). Body-mass index and incidence of cancer: a systematic review and meta-analysis of prospective observational studies. Lancet 371, 569–578.

Richardson, B., Price, A., Wagner, M., Williams, V., Lorigan, P., Browne, S., Miller, J. G. and Mac Neil, S. (1999). Investigation of female survival benefit in metastatic melanoma. Br. J. Cancer 80, 2025–2033.

Ringel, A. E., Drijvers, J. M., Baker, G. J., Catozzi, A., García-Cañaveras, J. C., Gassaway, B. M., Miller, B. C., Juneja, V. R., Nguyen, T. H., Joshi, S., et al. (2020). Obesity Shapes Metabolism in the Tumor Microenvironment to Suppress Anti-Tumor Immunity. Cell 183, 1848–1866 e26.

Sergentanis, T. N., Antoniadis, A. G., Gogas, H. J., Antonopoulos, C. N., Adami, H.-O., Ekbom, A. and Petridou, E. T. (2013). Obesity and risk of malignant melanoma: a meta-analysis of cohort and case-control studies. Eur. J. Cancer 49, 642–657.

Shutter, J. R., Graham, M., Kinsey, A. C., Scully, S., Lüthy, R. and Stark, K. L. (1997). Hypothalamic expression of ART, a novel gene related to agouti, is up-regulated in obese and diabetic mutant mice. Genes Dev. 11, 593–602.

Song, Y. and Cone, R. D. (2007). Creation of a genetic model of obesity in a teleost. FASEB J. 21, 2042–2049.

Song, Y., Golling, G., Thacker, T. L. and Cone, R. D. (2003). Agouti-related protein (AGRP) is conserved and regulated by metabolic state in the zebrafish, Danio rerio. Endocrine 22, 257– 265.

Tagore, M., Hergenreder, E., Suresh, S., Baron, M., Perlee, S., Melendez, S., Hollmann, T. J., Ideker, T., Studer, L. and White, R. (2021). Electrical activity between skin cells regulates melanoma initiation. BioRxiv.

Tung, Y. C. L., Rimmington, D., O’Rahilly, S. and Coll, A. P. (2007). Pro-opiomelanocortin modulates the thermogenic and physical activity responses to high-fat feeding and markedly influences dietary fat preference. Endocrinology 148, 5331–5338.

van der Klaauw, A. A., Keogh, J. M., Henning, E., Stephenson, C., Kelway, S., Trowse, V. M., Subramanian, N., O’Rahilly, S., Fletcher, P. C. and Farooqi, I. S. (2016). Divergent effects of central melanocortin signalling on fat and sucrose preference in humans. Nat. Commun. 7, 13055.

Verboven, K., Wouters, K., Gaens, K., Hansen, D., Bijnen, M., Wetzels, S., Stehouwer, C. D., Goossens, G. H., Schalkwijk, C. G., Blaak, E. E., et al. (2018). Abdominal subcutaneous and visceral adipocyte size, lipolysis and inflammation relate to insulin resistance in male obese humans. Sci. Rep. 8, 4677.

Weiss, J. M., Hunter, M. V., Tagore, M., Ma, Y., Misale, S., Simon-Vermot, T., Campbell, N. R., Newell, F., Wilmott, J. S., Johansson, P. A., et al. (2020). Anatomic position determines oncogenic specificity in melanoma. BioRxiv.

Wolk, A., Gridley, G., Svensson, M., Nyrén, O., McLaughlin, J. K., Fraumeni, J. F. and Adam, H. O. (2001). A prospective study of obesity and cancer risk (Sweden). Cancer Causes Control 12, 13–21.

Zang, L., Maddison, L. A. and Chen, W. (2018). Zebrafish as a model for obesity and diabetes. Front. Cell Dev. Biol. 6, 91.

Zhang, L., Alt, C., Li, P., White, R. M., Zon, L. I., Wei, X. and Lin, C. P. (2012). An optical platform for cell tracking in adult zebrafish. Cytometry A 81, 176–182.

Zhang, M., Di Martino, J. S., Bowman, R. L., Campbell, N. R., Baksh, S. C., Simon-Vermot, T., Kim, I. S., Haldeman, P., Mondal, C., Yong-Gonzales, V., et al. (2018). Adipocyte-Derived Lipids Mediate Melanoma Progression via FATP Proteins. Cancer Discov. 8, 1006– 1025.

